# De Novo sequencing-assisted homology search for DIA data analysis enables low abundance peptide variants discovery

**DOI:** 10.1101/2025.05.30.657054

**Authors:** Rui Qiao, Hechen Li, Haibo Bian, Lei Xin, Baozhen Shan

## Abstract

Data-independent acquisition mass spectrometry (DIA-MS) has emerged as a powerful approach for comprehensive proteome profiling. Spectral library search and library-free search are the two major approaches for DIA data analysis. The spectral library search requires high-quality spectral libraries derived from the search results of data-dependent acquisition (DDA) experiments, while library-free approaches rely on prediction models to generate in silico libraries. Both methodologies constrain the search space to the peptide list in the database, limiting the discovery of variant peptides arising from genetic variations or mutations. We present a novel computational method DIAVariant designed to identify peptide sequence variants directly and solely from complex DIA spectra while rigorously controlling the false discovery rate. Our experimental results demonstrate that DIAVariant successfully identifies sequence variants previously detected through proteogenomic approaches, while maintaining high specificity across multiple datasets. When integrated with existing DIA database search solutions, our approach constitutes a comprehensive analytical workflow capable of identifying peptides both represented within reference protein databases and those arising from sequence variations not captured in standard databases.

## 1 Introduction

Data-independent acquisition mass spectrometry (DIA-MS) has revolutionized the field of proteomics, rapidly gaining prominence as a powerful and comprehensive technique for protein identification and quantification. Unlike traditional data-dependent acquisition (DDA) methods, which selectively fragment and analyse the most abundant precursor ions in a sample, DIA-MS systematically fragments all ions within a defined mass range during each scan cycle. This comprehensive fragmentation approach offers several advantages over DDA, leading to its increasing adoption in the proteomics community. Specifically, DIA-MS provides improved quantitative accuracy, reproducibility, and higher proteome coverage [1, 2].

The increasing popularity of DIA-MS can be attributed to several key factors. Firstly, its ability to generate more complete and reproducible quantitative data allows for more reliable comparison of protein expression levels across different samples or conditions. Secondly, the inherent nature of DIA, where all fragment ions are recorded, allows for deeper proteome coverage than DDA. This is because low-abundance peptides, which might be missed by the selective nature of DDA, are more likely to be identified and quantified in a DIA experiment. Finally, the development of sophisticated data analysis tools and workflows has made the processing and interpretation of DIA data more accessible [3, 4, 5], further driving its widespread adoption.

Sequence variants in proteins, arising from single nucleotide polymorphisms (SNPs), insertions, deletions, and other genetic mutations, play a crucial role in various biological processes and disease. These variants can alter protein structure, function, stability, and interactions, potentially leading to phenotypic differences, drug resistance, or disease susceptibility [6, 7, 8, 9]. Identifying sequence variants at the protein level is therefore critical for understanding the molecular basis of human disease, as well as for developing personalized medicine strategies.

Despite the inherent advantages of DIA-MS, the identification of sequence variants from DIA data presents substantial methodological and computational challenges. Current DIA analysis methodologies predominantly rely on spectral matching against libraries derived from DDA experiments or in silico predictions based on genomic databases [3, 4]. This library-dependent paradigm inherently constrains the discovery of novel peptides absent from the reference libraries, particularly those harbouring sequence variants. While proteogenomic approaches that integrate genomic and proteomic data can facilitate the construction of comprehensive variant-aware libraries [10], these methodologies typically require specialized data acquisition protocols and sequential data analysis pipelines which might carry mistakes from upstream genomics data over to downstream proteomics data analysis. Moreover, recent investigations in neoantigens have revealed that sequence variants may originate from alternative splicing events and previously unannotated non-coding regions [11, 12], which lie beyond the scope of conventional proteogenomic approaches. Recently, various de novo sequencing methodologies have been proposed for novel peptide identification from DIA data [13, 14]. These approaches frequently employ complex neural network architectures that function as black box models and the predicted sequences lack interpretability. Furthermore, their validation has been predominantly limited to sequences present in existing libraries, leaving considerable uncertainty regarding the false discovery rate among the reported novel peptides. In this work, we present a novel computational framework DIAVariant for the direct identification of sequence variants from DIA-MS data without reliance on prior genomic information. Our approach extends conventional library-dependent methodologies by implementing a two-stage analysis pipeline. First, we leverage the PEAKS DIA algorithm [15], which facilitates the identification of both library-annotated and novel peptides from complex DIA spectra. Subsequently, we employ the SPIDER search algorithm [16] to correct de novo sequencing errors through reference genome alignment, thereby identifying peptide sequences that diverge from reference peptides by only one or two amino acid substitutions. To enhance the specificity of our variant identification, we implement a secondary validation process wherein we predict the indexed Retention Time (iRT) values and theoretical fragment ion spectra for putative variant sequences. These predictions enable a second round of library searching, effectively filtering out novel sequence precursors that exhibit excessive fragment ion signal overlap with confidently identified reference peptides. This two-stage search strategy will give us a list of novel peptide variants supported by evidence from both liquid chromatography (LC) and MS dimensions, accompanied by statistically meaningful scores that facilitate stringent false discovery rate control. Our experimental validation demonstrates that the proposed methodology successfully identifies sequence variants previously detectable only through proteogenomic approaches.

## 2 Methods

Our workflow is peptide-centric and begins with MS1 feature detection to identify reliable MS1 signals with well-defined isotopic patterns characteristic of ionized peptides.

Subsequently, we perform PEAKS DIA database search against the user-provided reference protein database. Precursors identified at 1% FDR are then mapped to previously detected MS1 features based on retention time and precursor m/z correspondence.

For MS1 features lacking confident peptide identification from the reference database, we implement DIA de novo sequencing [13] to predict peptide sequences that optimally match the MS2 scans associated with each original MS1 feature. The SPIDER algorithm [16] is then applied to confident de novo candidates, utilizing the de novo peptide sequence, MS1 feature information, and reference database to correct potential de novo sequencing errors and identify sequences that could align to the reference protein database.

All SPIDER candidate peptides undergo an initial filtering, retaining only peptides with one or two mutations. We then predict the indexed retention time (iRT) for these potential mutated peptides and convert it to experimental retention time using the RT regression model established during the previous database search. Peptides exhibiting substantial discrepancies between predicted and observed retention times are eliminated from consideration.

To control the quality of the remaining sequence variants, we run another round of spectral library search. This library is constructed on-the-fly by combining the remaining candidate peptide variants with peptides identified in the preceding DIA database search. Retention times in the library are assigned according to the empirical values from previous search results, and a PEAKS DIA spectral library search specific [15] to the data is conducted by directly targeting these retention times. Q-values are reported for each precursor, allowing users to apply appropriate thresholds and manually evaluate the biological significance of reported sequence variants.

For any given MS sample, our workflow reports two sets of identification results. One is the set of sequences from the protein database, with an FDR estimate using the traditional target-decoy search strategy. The other one is a set of sequence variants that might exist in the original biological sample. A flow chart of DIAVariant is demonstrated in Figure 1.

**Figure 1.**
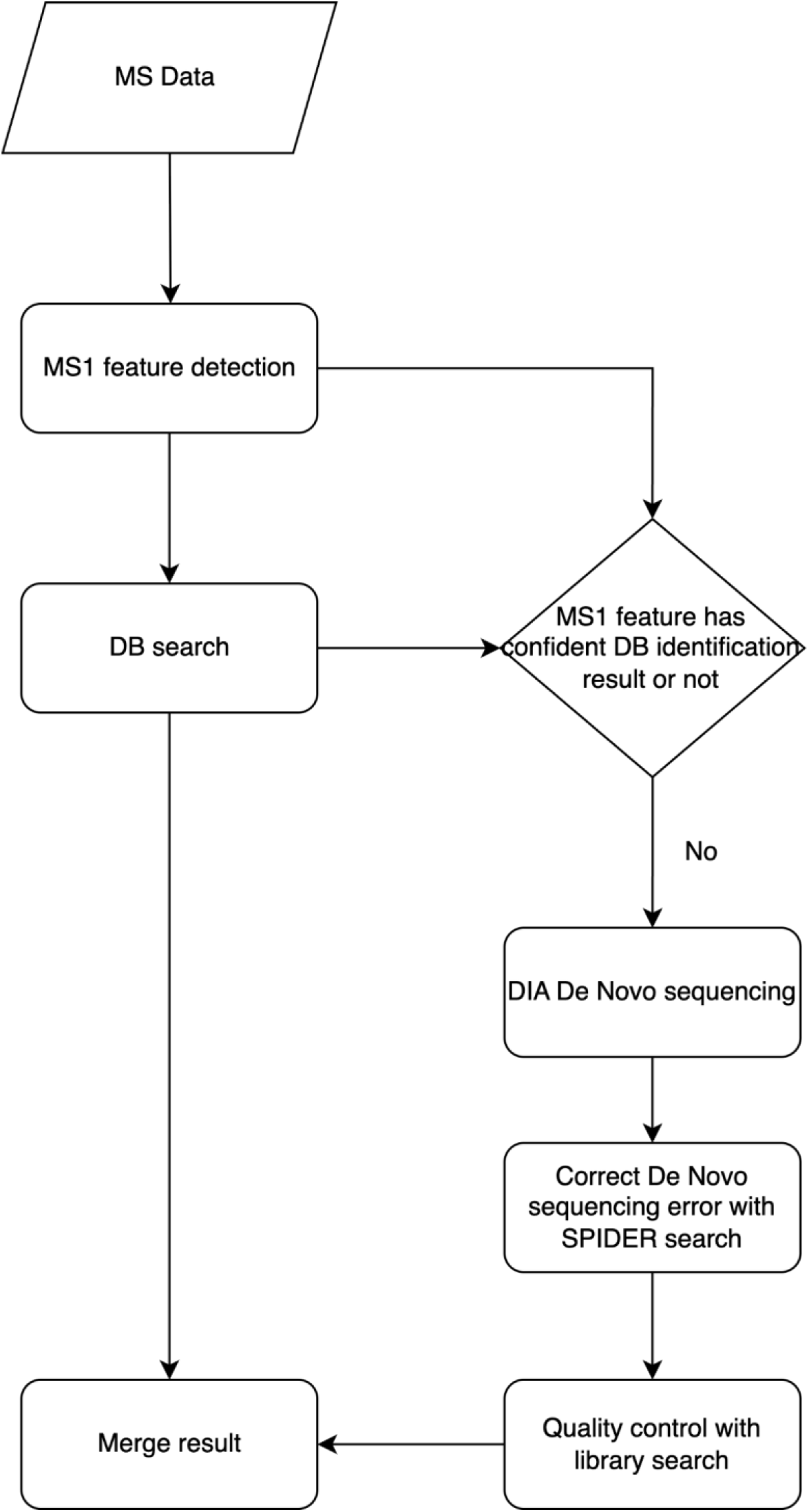
DIAVariant workflow

## 3 Results

As illustrated in Figure 1, DIA DB search is integrated within our proposed analytical workflow. The implementation of accurate and sensitive DB search methodologies is essential for minimizing false positive identifications in sequence variants. To validate the performance of our DIA DB search engine, we benchmarked PEAKS DIA DB against two recently published datasets [17][18]. Our results in Table 1 demonstrates that the PEAKS DIA DB search algorithm achieves identification and quantification performance metrics that are at least comparable to those obtained with existing DIA solutions.

**Table 1.**
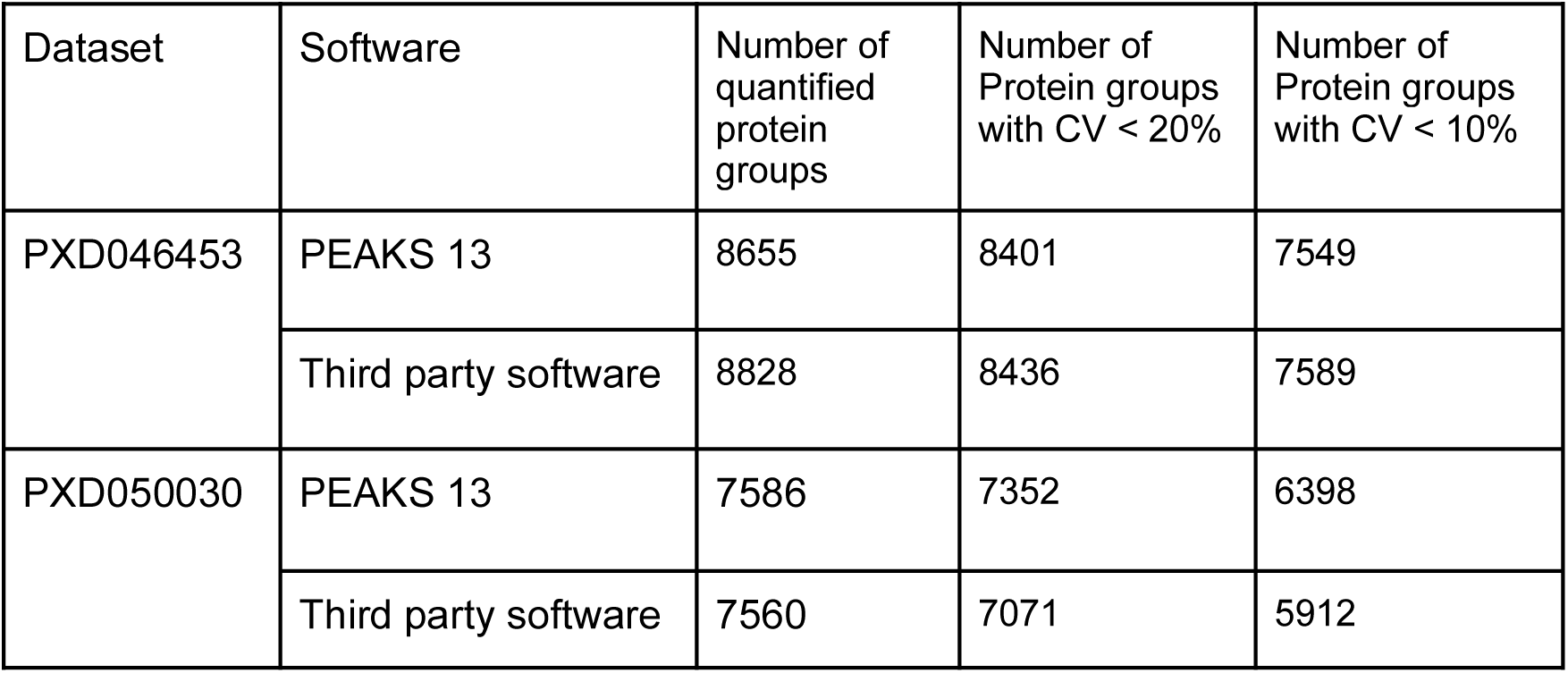
PEAKS DIA DB benchmarks.

To assess the specificity of sequence variants identified by our workflow, we conducted validation using public MS datasets analysed against non-native reference protein databases from homologous species. For instance, human samples were searched against a mouse protein database. This cross-species validation approach enabled us to evaluate the biological relevance of reported sequence variants by determining the proportion that could be confidently identified (i.e. q-value < 0.01) when searched against the original species’ protein database. We conducted the validation on ABRF data [19] (human sample, search against mouse DB) and a recent dataset from Orbitrap Astral [20] (MSV000095360, mouse sample, search against human DB). As demonstrated in Table 2, more than 85% of the peptide variants reported by DIAVariant are confident precursors from the native reference database.

**Table 2.** Cross-species validation result.

|  | Number of reported sequence variants | Number of sequence variants that can be confidently identified in the native database | Accuracy |
| --- | --- | --- | --- |
| ABRF data | 340 | 315 | 92.65% |
| MSV000095360 | 964 | 844 | 87.55% |

We then applied DIAVariant to analyse the dataset published by Fierro-Monti et al. [10]. In their investigation, Fierro-Monti et al. performed exome sequencing on HeLa cell line samples and conducted variant calling to identify 233 canonical HeLa variant protein sequences. These variant sequences were then incorporated into the human reference protein database to generate a customized protein database. The authors then searched DIA MS spectra from the HeLa samples against this customized database using DIA-NN 1.8.1. Among the sequence variants identified at 1% FDR, Fierro-Monti et al. reported six peptides validated through stable isotope dilution and parallel reaction monitoring. In contrast to the proteogenomic approach employed by Fierro-Monti et al., our proposed method operates directly and exclusively from MS data, which means confident MS1 precursor isotope envelopes are needed to confirm the presence of peptide variants in the original sample. Upon thorough examination of the raw MS scans, we determined that four of the six previously reported variant peptides lacked sufficient MS1 signals and consequently could not be detected by our method. Notably, for the remaining two peptides with adequate MS1 evidence, our method successfully retrieved them on several DIA MS samples. Table 3 shows a summary of this result. The matching spectra of the two peptides are demonstrated in Supplementary Figure 1-7. In addition to the two previously reported variant peptides, DIAVariant identified 426 additional variants. Several of these variants exhibited low abundance, characterized by low MS1 precursor intensity, and consequently were hard to be detected in DDA data. Supplementary Figure 8 presents a representative example of such low-abundance variants detected by DIAVariant.

**Table 3.** Our method finds 2 out of the 6 previous reported sequence variants.

| Peptide variants reported by proteogenomic method | Our proposed method |
| --- | --- |
| LEQDLQQIAK | Found in 2/5 samples |
| ERLEQDLQQIAK | No MS1 signal |
| QGTfHSQQALEYGTR | No MS1 signal |
| KIEGDLQGDHqK | No MS1 signal |
| VYVVPmGGVR | No MS1 signal |
| NELSGALTGLIR | Found in 5/5 samples |

## 4 Discussion

In this study, we present a novel workflow for direct identification of sequence variants exclusively from DIA MS data. Our experimental results demonstrate that DIAVariant successfully identifies previously reported sequence variants detected through proteogenomic approaches, given the presence of robust MS1 signals. In addition, DIAVariants reported hundreds more peptide variants of different abundance. The biological validation of these candidate variants requires further experimental investigation and represents an important avenue for future research.

Notably, our approach exhibits potentials for detecting sequence variants arising from alternative splicing events or non-coding genomic regions—variants that remain largely inaccessible to conventional proteogenomic methodologies. This represents a substantial advancement in variant detection capabilities within proteomics research, addressing critical gaps in existing analytical frameworks.

## Supplementary information

**Supplementary Figure 1.**
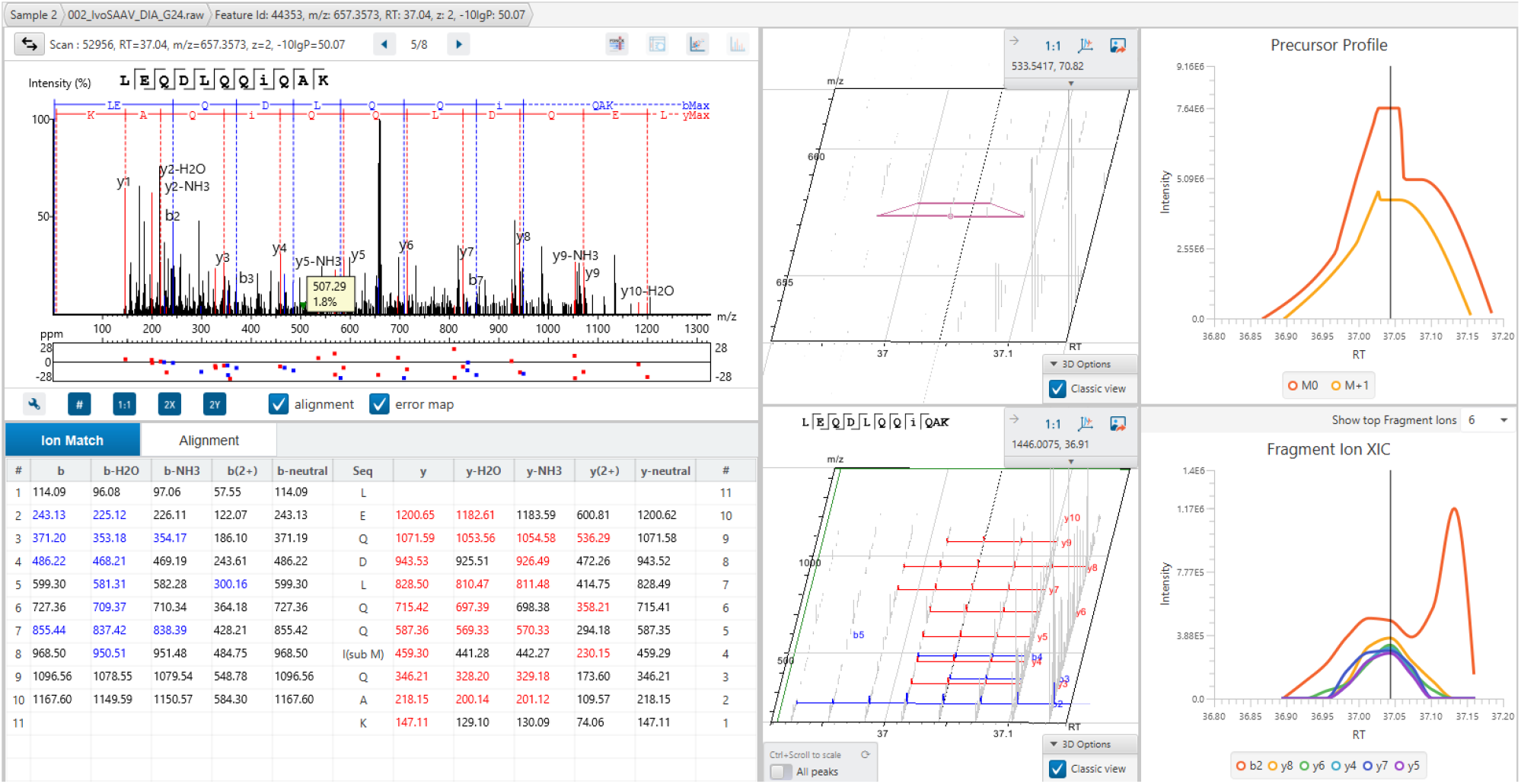
LEQDLQQIQAK in Sample 2

**Supplementary Figure 2.**
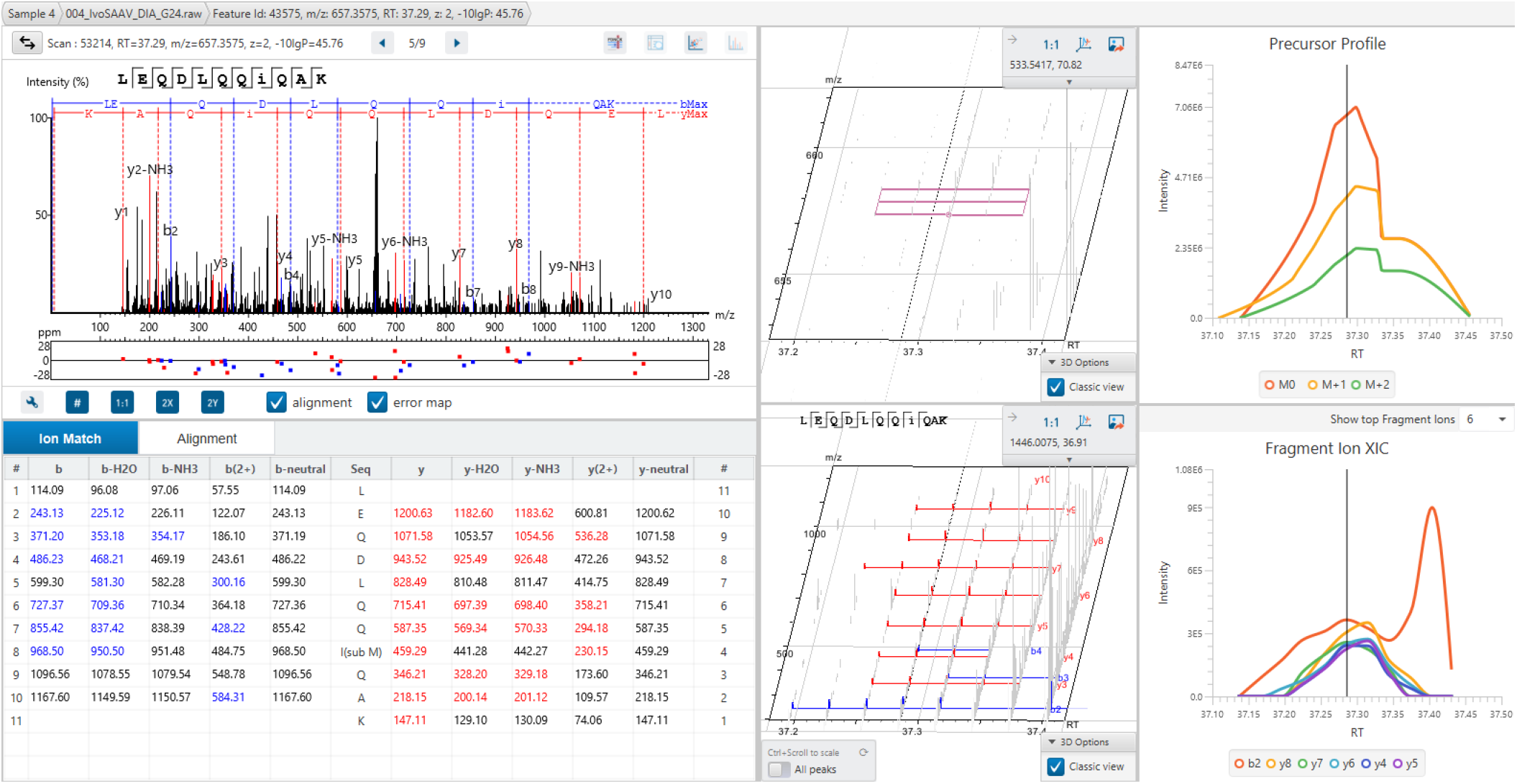
LEQDLQQIQAK in Sample 4

**Supplementary Figure 3.**
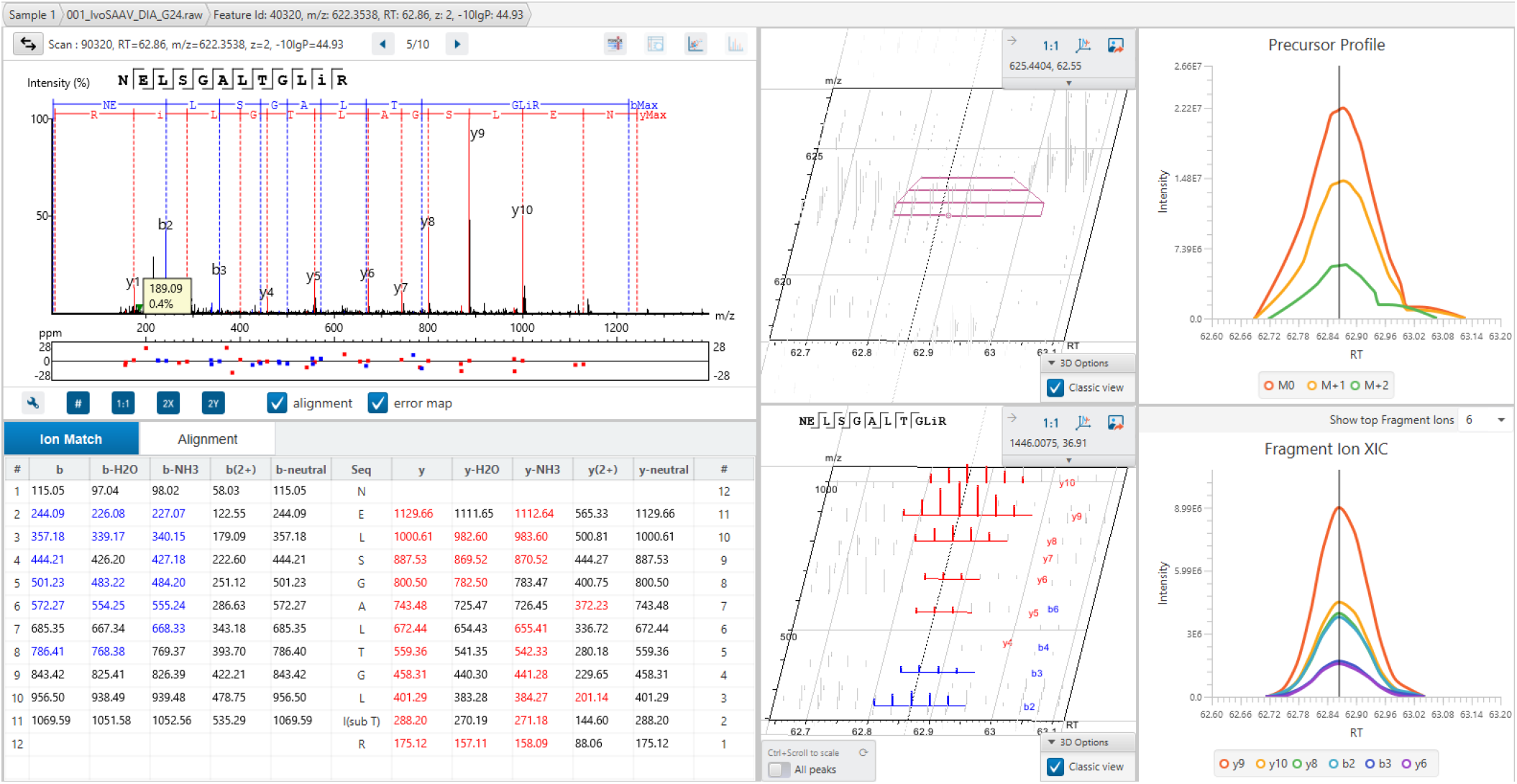
NELSGALTGLIR in Sample 1

**Supplementary Figure 4.**
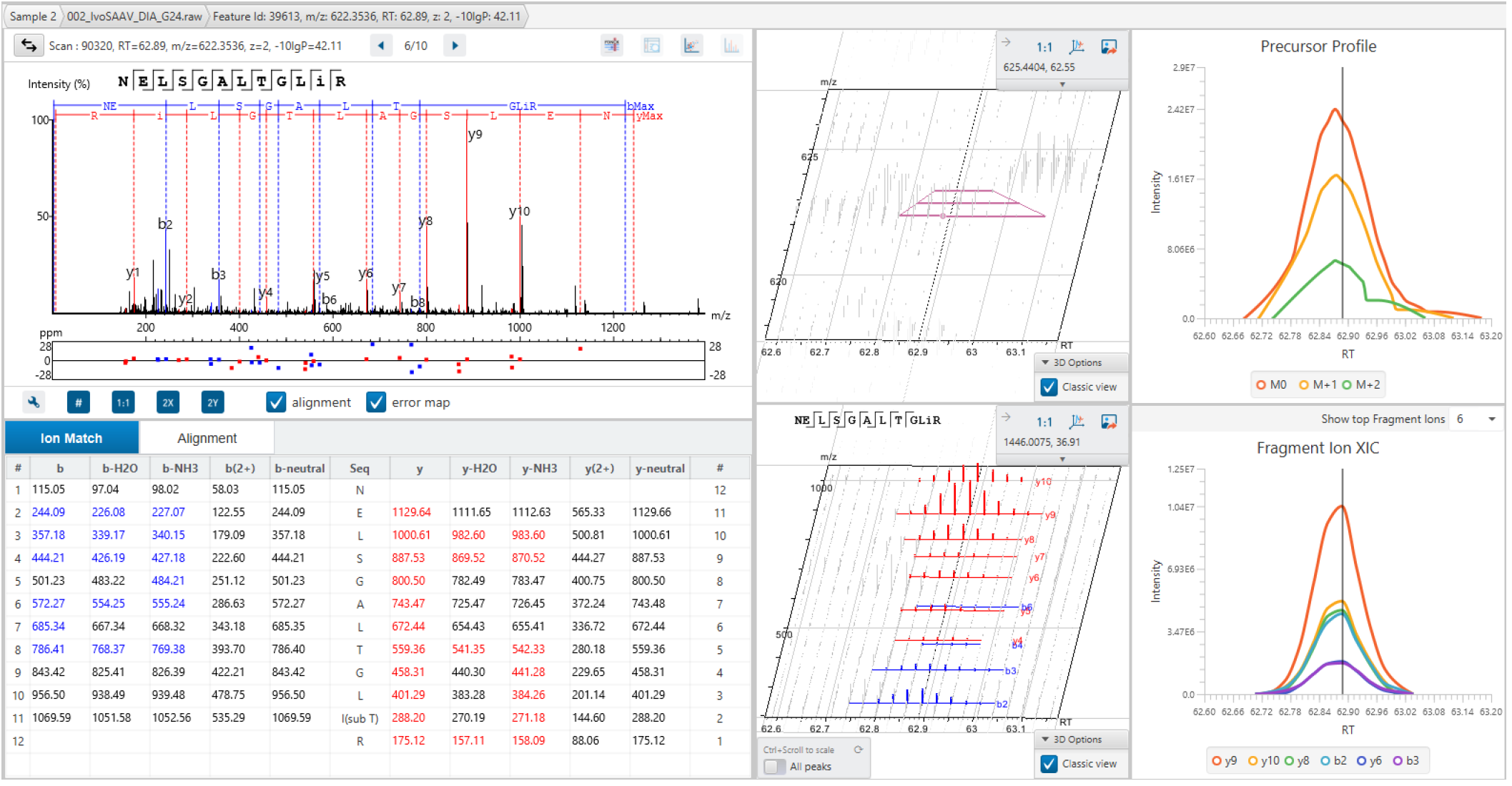
NELSGALTGLIR in Sample 2

**Supplementary Figure 5.**
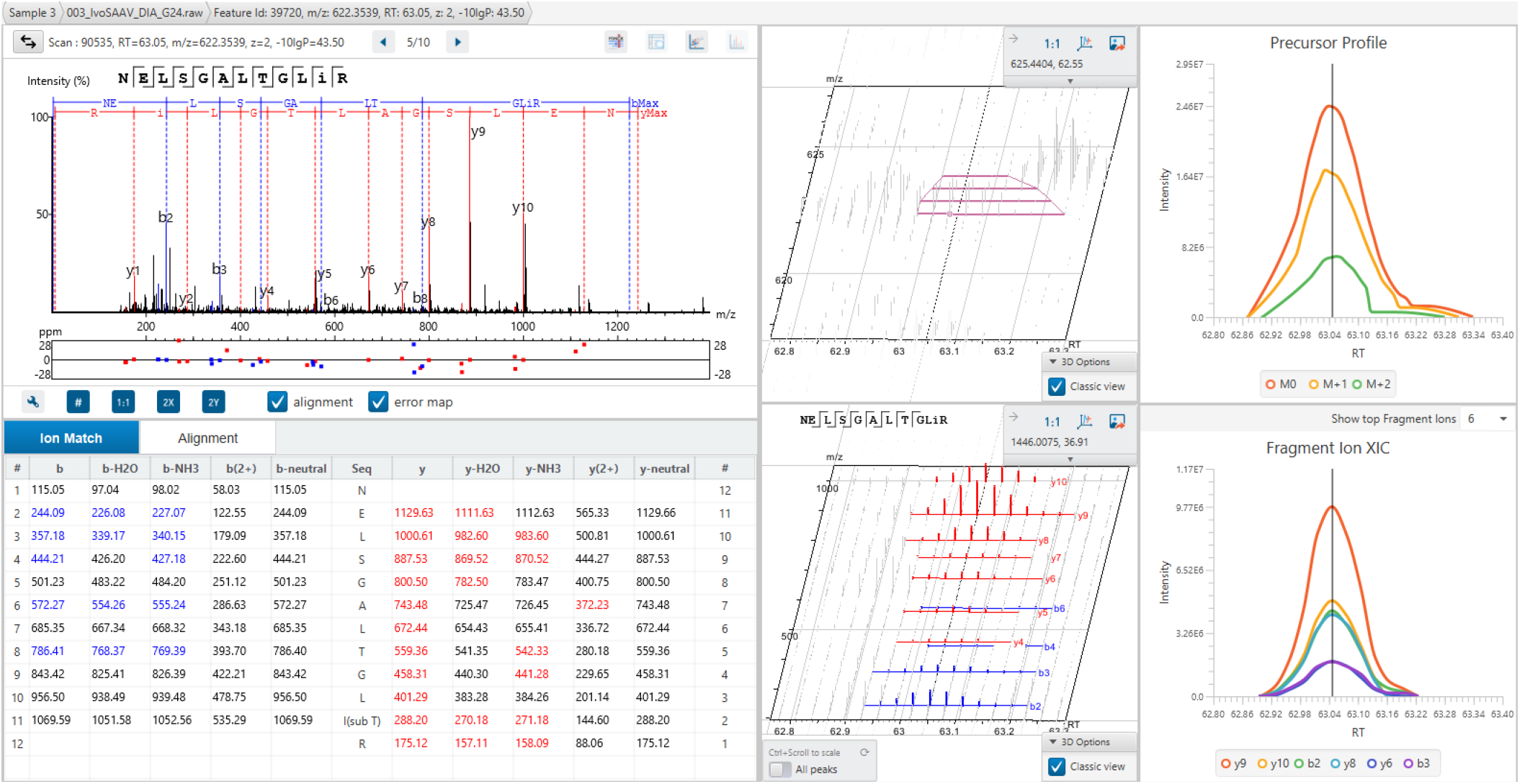
NELSGALTGLIR in Sample 3

**Supplementary Figure 6.**
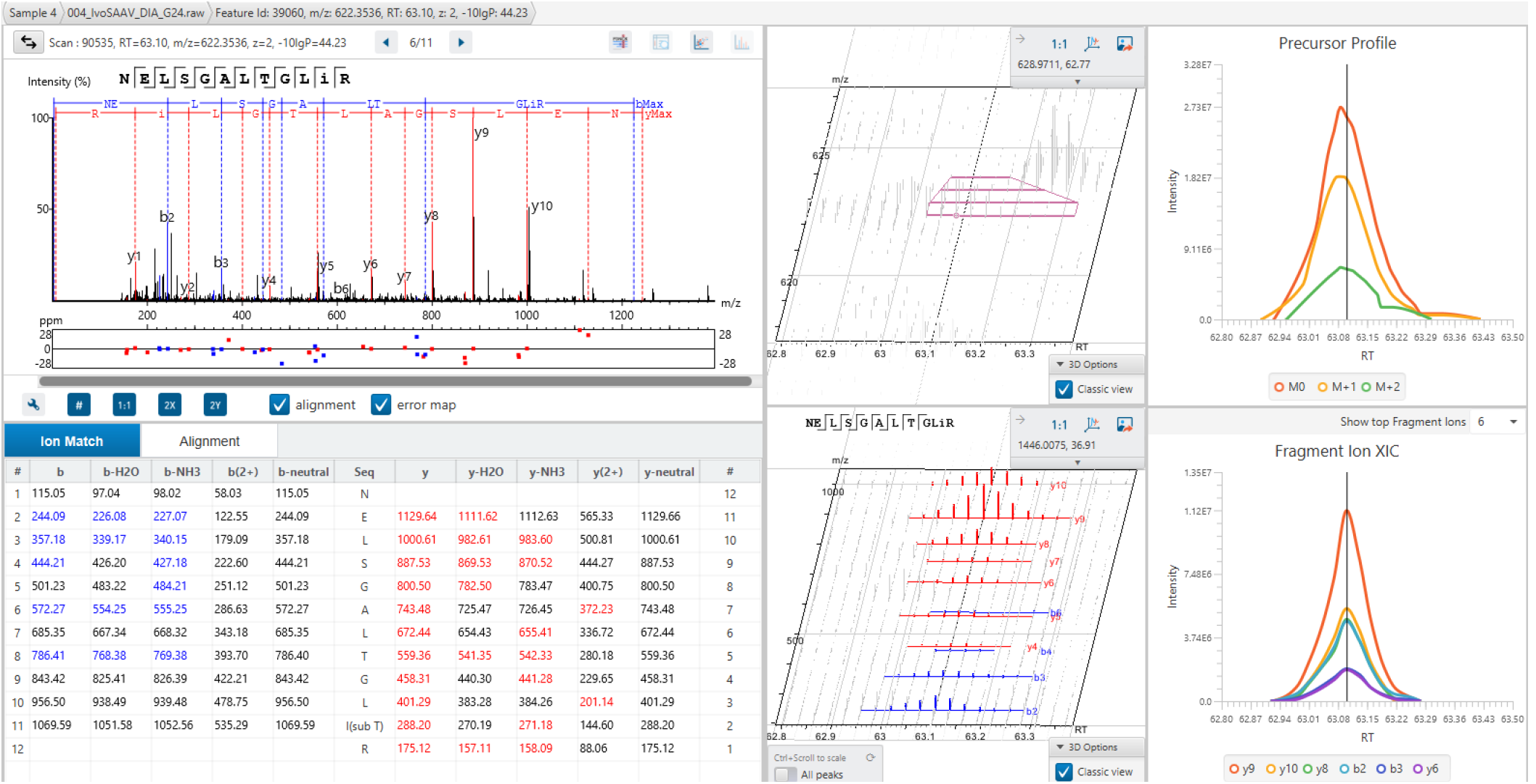
NELSGALTGLIR in Sample 4

**Supplementary Figure 7.**
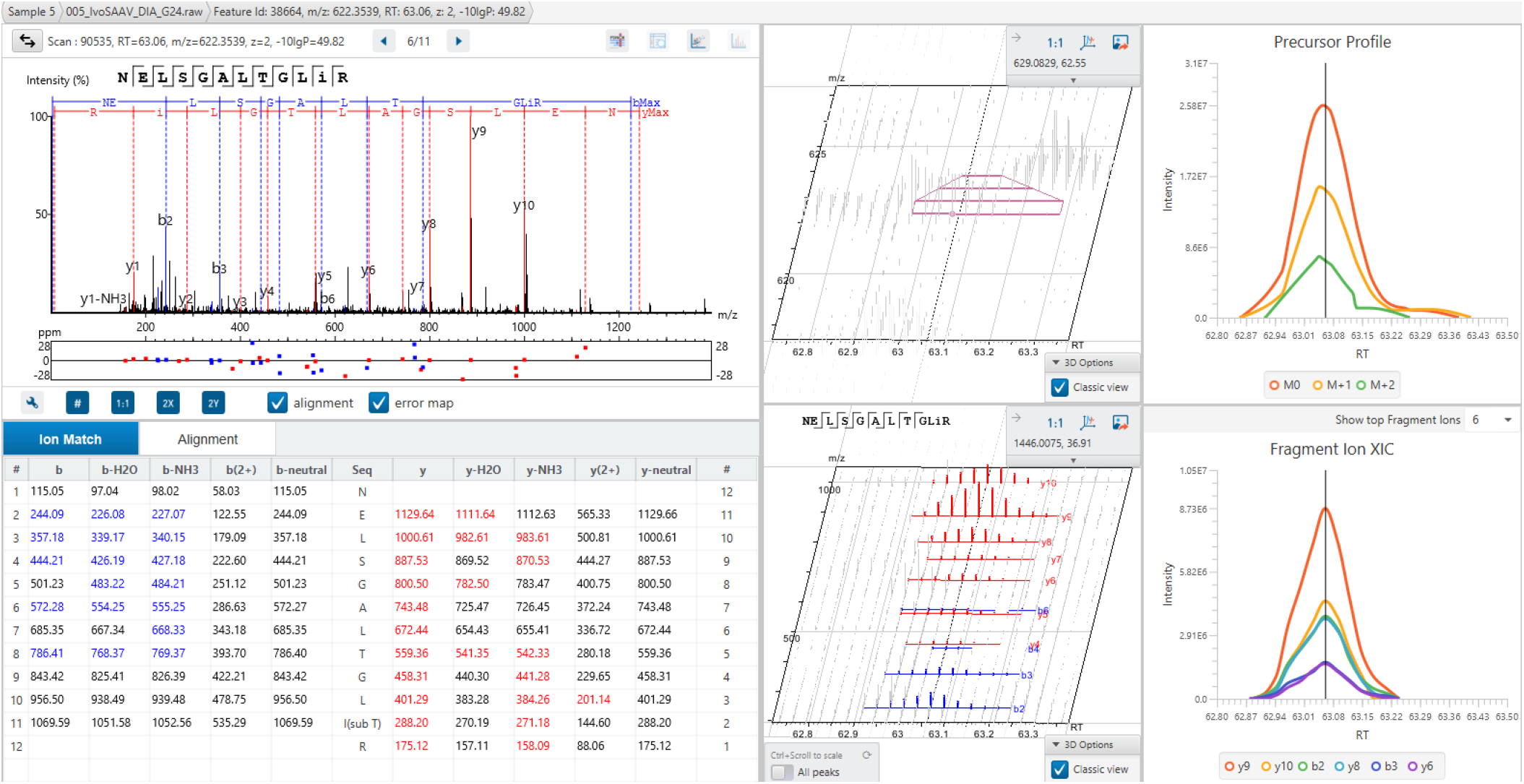
NELSGALTGLIR in Sample 5

**Supplementary Figure 8.**
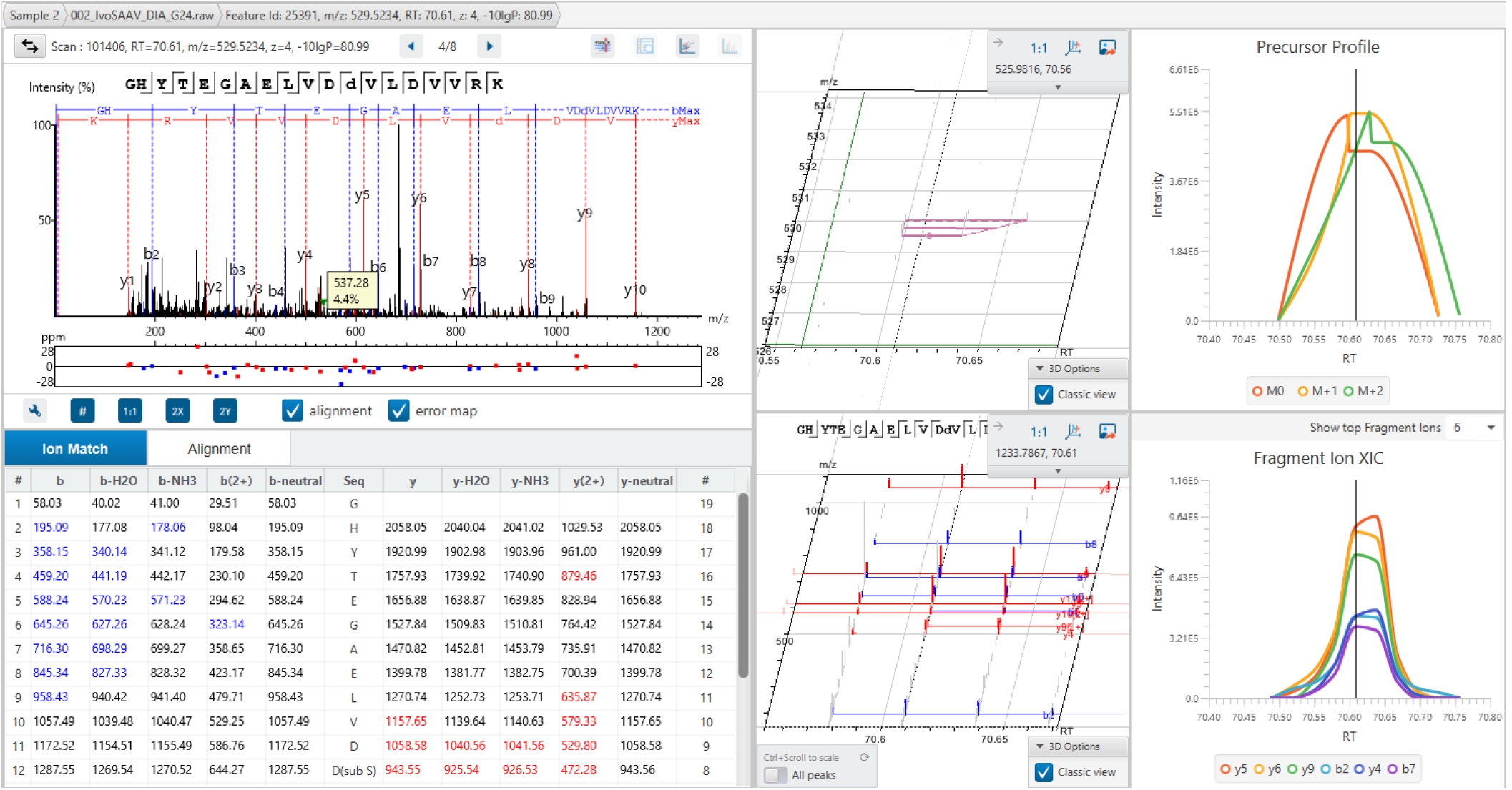
A low abundant peptide variant reported by DIAVariant.

